# RUSBoost: A suitable species distribution method for imbalanced records of presence and absence. A case study of twenty-five species of Iberian bats

**DOI:** 10.1101/2021.10.06.463434

**Authors:** Jaime Carrasco, Fulgencio Lisón, Andrés Weintraub

## Abstract

1. Traditional Species Distribution Models (SDMs) may not be appropriate when examples of one class (e.g. absence or pseudo-absences) greatly outnumber examples of the other class (e.g. presences or observations), because they tend to favor the learning of observations more frequently.
2. We present an ensemble method called **R**andom **U**nder**S**ampling and **Boost**ing (RUSBoost), which was designed to address the case where the number of presence and absence records are imbalanced, and we opened the “black-box” of the algorithm to interpret its results and applicability in ecology.
3. We applied our methodology to a case study of twenty-five species of bats from the Iberian Peninsula and we build a RUSBoost model for each species. Furthermore, in order to improve to build tighter models, we optimized their hyperparameters using Bayesian Optimization. In particular, we implemented a objective function that represents the cross-validation loss: 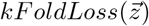, with 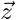 representing the hyper-parameters *Maximum Number of Splits, Number of Learners* and *Learning Rate*.
4. The models reached average values for Area Under the ROC Curve (AUC), specificity, sensitivity, and overall accuracy of 0.84 ± 0.05%, 79.5 ± 4.87%, 74.9 ± 6.05%, and 78.8 ± 5.0%, respectively. We also obtained values of variable importance and we analyzed the relationships between explanatory variables and bat presence probability.
5. The results of our study showed that RUSBoost could be a useful tool to develop SDMs with good performance when the presence/absence databases are imbalanced. The application of this algorithm could improve the prediction of SDMs and help in conservation biology and management.

## 1 Introduction

Artificial intelligence methods, particularly those of Machine Learning (ML), are useful in finding complex interactions between the studied variables directly from the data (Flach, 2001; Olden et al., 2008), and have been recognized as formidable scientific tool in natural resource management and wildlife biology (Elith and Leathwick, 2009; Olden et al., 2008). ML approach can be applied to virtually any data and eliminates the need to specify, *a priori*, broadly untested and potentially biased assumptions regarding the underlying statistical distribution of the data (Breiman et al., 2001). Moreover, various studies have indicated that traditional statistical methods are not suitable for problems that come from ecology or related fields due to their low predictive power (Breiman et al., 2001; Elith et al., 2008).

Most of the tools for modeling species distribution today come from ML methods. In particular, supervised learning models are closely related to Species Distribution Models (SDMs), which attempts to predict the probability of the presence of a species, given certain variables that characterize the underlying environment (Wisz et al., 2013; Jiménez-Valverde et al., 2013; Thibaud et al., 2014; García-Callejas and Araújo, 2016). In the last decade, several methods such as Classification and Regression Trees (Breiman et al., 2017), Random Forest (Breiman, 2001), Boosted Trees (Freund et al., 1996), Support Vector Machines (Schölkopf et al., 2001), Artificial Neural Networks (Bishop et al., 1995; Goodfellow et al., 2016) and so forth, have been intensively used in different applications that involve the modeling of the species distribution, or also commonly referred to as ecological niche models (ENMs) (Cutler et al., 2007; De’Ath, 2007; Drake et al., 2006; Lek and Guégan, 2012; Olden et al., 2008; Phillips et al., 2006). In general terms, the purpose of the SDM is to predict the spatial variation of the presence or abundance of species within the occupied environmental space. For this purpose, biotic or abiotic variables/drivers are constructed, usually associated with the geographical place where the species has been observed, and depending on the modeling approach, variables are also assigned to the places where they have not been documented (Elith and Leathwick, 2009; Pulliam, 2000; Soberón and Nakamura, 2009). Based on the information provided in the modeling process, SDMs can be divided into two branches: i) those that are based on data that contain only presence (Elith et al., 2006; Thibaud et al., 2014; Wisz et al., 2013; Tsoar et al., 2007) or, ii) those that record the presences and absences (De’Ath, 2007; Cutler et al., 2007). In ML, the second branch is a typical two-class classification problem, or also called a binary classification (presence and absence classes). In contrast, the former has been developed mainly using methods that do not come directly from the ML approach, for example Convex Hull approach (Cornwell et al., 2006) or BIOCLIM using depth functions (Busby, 1991), to name a few examples.

There is still a debate about which of the two approaches is the most appropriate. Some comparative studies indicated that the models that used presence and absence data were superior to those that only used presence (Brotons et al., 2004; Elith et al., 2006). Others emphasize presence-only models for conceptual reasons, stating that one-class models more elegantly represent the idea of niche rather two-class models (Drake et al., 2006; Drake and Bossenbroek, 2009); and some recommend one or the other approach depending: if the absence data is reliable, or if it is necessary to optimize the sensitivity or specificity metrics (Maher et al., 2014). However, what is not up for discussion is the balancing of data in the presence/absence approach: whether certain classes, for example the presence observations are underrepresented in the dataset, traditional models tend to favor the learning of observations more frequently, producing a classifier biased towards the majority class (Richhariya and Tanveer, 2020; Seiffert et al., 2009). For example, when fauna is modeled, it is common that this phenomenon is likely to occur, due to the biological and behavioral characteristics of the animals (elusive animals, misidentification in the capture processes, incomplete or biased monitoring, inaccessible areas, rare species, etc.) (Soley-Guardia et al. (2016); Sillero et al. (2021); Velasco and González-Salazar (2019)). Therefore, finding algorithms capable of improving predictions with this type of imbalance is important when it comes to providing useful tools for managing biodiversity conservation at different levels.

The problem of learning from imbalanced data has been carefully dealt in different ways and for different methods. (Chawla et al., 2002) introduced a method that over-samples the minority class and under-samples the majority class called Synthetic Minority Oversampling Technique (SMOTE), and tested its solution with an algorithm used to generate a decision tree (C4.5) (Quinlan, 1992), a Rule-based classification algorithm called Ripper (Cohen, 1995) and a Naive Bayes classifiers (Mladenic and Grobelnik, 1999). Seiffert et al. (2009) combined Random UnderSampling (RUS) and Boosting (RUSBoost) (Freund et al., 1996) providing a simple and efficient method for improving predictive performances when the database containing the species’ presence and absence records were imbalanced. Recently, Richhariya and Tanveer (2020) uses oversampling and undersampling of data to remove the imbalance in the classes and “*universum*” learning with SVM method. Currently, these techniques and procedures that deal with the problem of data imbalance have been little explored in studies whose purpose is modeling the distribution of species and only there are few in the literature (Breiner et al. (2015), Johnson et al. (2012), and Robinson et al. (2018)).This is an important issue when the data comes from species which are problematic in their identification or which have not been subjected to an equal survey effort.

In our study, we hypothesized that the RUSBoost method is suitable for modeling the distribution of species that contain records of presence and absence data, and its performance is independent of the level of imbalance between presence and absence data. Therefore, our aims were: i) to explain in detail how the method works, in algorithmic and interpretability and applicability in ecological terms; and, ii) to apply our methodology to a case study of twenty-five species of bats from the Iberian Peninsula and analyze the results. In addition, we present the Bayesian Optimization procedure (Eggensperger et al., 2013) to find the optimal hyperparameters of the method, to achieve the best performance of the models (one for each species).

## 2 Materials and methods

### 2.1 Context, bat species and predictors

Bats are a diverse mammals group very threatened and they need conservation policies. However, bats are nocturnal animals and the different survey techniques could affect our capacity to detect some species. Therefore, it is usual to find distribution maps imbalanced for these mammals. For this reason, we used presence/absence distribution data of bat species present in the Iberian Peninsula. This is a biogeographical region with an area about six millions of square kilometers divided into 10 × 10 *km*^2^ UTM cells. The dataset was elaborated from the distribution maps of “Atlas y Libro Rojo de los Mamíferos Terrestres de España” (Palomo et al., 2007) and “Atlas dos morcegos de Portugal” (Rainho et al.); and was recently updated with new records with the same resolution form personal surveys and references (Lisón et al., 2015; Lisón and Sánchez-Fernández, 2017). We did not use those bat species where there were less that thirty records. Also, we did not include the twin species *Myotis nattereri/escalerai* because there is uncertainty with its distribution (Palomo et al., 2007; Juste et al., 2018). For the species *Eptesicus serotinus* and *E. isabellinus*, we considered the distribution of the last one as limited to south and south eastern of Iberia (Andalusia and Murcia) (Palomo et al., 2007; Lisón et al., 2015).

Land-use variables were obtained following the procedure by Lisón et al. (2015) and Lisón and Sánchez-Fernández (2017). We used the land-use map provided by the CORINE Land Cover Project (see www.eea.europa.eu). Also, we modelled the habitat suitable models of bat species using as environmental variables the land-use cover from CORINE Land Cover Project for 2006 (https://land.copernicus.eu/pan-european/corine-land-cover/clc-2006). The land-use cover was reclassified from the 44 initial categories to 16 final categories (see Lisón and Sánchez-Fernández (2017)). For each final category, we calculated the percentage of surface occupied in each cell. Additionally, we used 44 environmental and geographical variables extracted of BIOMOD for each cell following Barbosa et al. (2012) (Table 1).

**Table 1:** Description of predictor variables.

| Drivers | Variable | Description | Variable | Description |
| --- | --- | --- | --- | --- |
| Human Activity | u100 | Distance to the nearest town with more than 100,000 inhabitants (km) | u500 | Distance to the nearest town with more than 500,000 inhabitants (km) |
|  | daut | Minimum distance road (km) | - | - |
| Geographic and Topographic | alti | Average altitude (m) | dalt | Difference in altitude (slope, m) |
|  | pend | Average slope | lati | Latitude |
|  | long | Longitude | - | - |
| Weather | tene | Average temperature in January | huen | Average humidity in January |
|  | tjul | Average temperature in July | huju | Relative humidity in July |
|  | tmed | Average annual temperature (°C) | vhum | Annual relative air humidity range |
|  | vtem | Annual temperature range (°C) = (TJul-TJan) | perm | Soil Permeability |
|  | inso | Mean annual insolation (hours/year) | rads | Mean annual sun radiation (Kwh/m<sup>2</sup>/day) |
|  | etp | Mean annual potential evapotranspiration (mm) | etr | Mean annual real evapotranspiration |
|  | prec | Mean annual precipitation (mm) | dipr | Mean annual number of days with precipitation >= 0.1 mm |
|  | pm24 | Maximum precipitation in 24 hours | pmr | Relative maximum precipitation (=PM24/Pmed) |
|  | dihe | Frost days | - | - |
| Land cover | Improductive | Improductive [%] | Urban | Urban areas [%] |
|  | Roads | Road [%] | Mines | Mine zones [%] |
|  | Rainfallcrops | Rainfall crops [%] | Irrigatedcrops | Irrigate crops [%] |
|  | Pasture | Pasture [%] | Mixedcrops | Mixed crops [%] |
|  | Abandoned | Abandoned crops [%] | Agroforestry | Agroforestry areas [%] |
|  | Forest | Forest [%] | Shurbland | Shrubland |
|  | SandRock | Sand and rock [%] | River | Rivers [%] |
|  | Wetland | Wetland [%] | - | - |

Our database was characterized by the high degree of imbalance in the records of presence and absence among bat species and geographic range that the species cover. Fig. 1 shows that the number of records comprise a range from 33 to 2, 297, with quartiles of *Q*_1_ = 212.5 and *Q*_2_ = 522 and *Q*_3_ = 914.25. Furthermore, the records were very scattered covering a latitudinal range of 36.06° to 43.7° (*Q*_1_ = 39.42, *Q*_2_ = 41.05, *Q*_3_ = 42.21) and longitudinal range of − 9.47° to 3.29° (*Q*_1_ = − 6.53, *Q*_2_ = − 4.46, *Q*_3_ = − 2.06).

**Figure 1:**
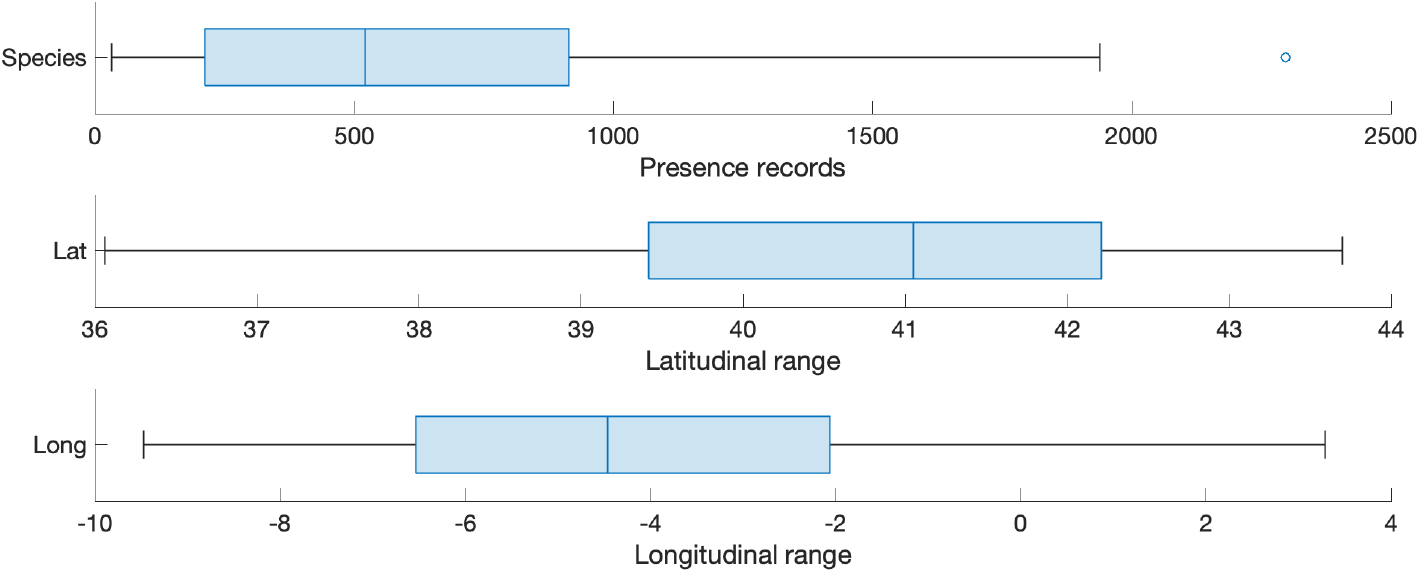
Bat records.

The basic notation and terminology that will be used throughout this section will be as follows: let 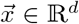 be the predictor variables described in Table 1 (then *d* = 36), and by *y* ∈ {0, 1} response variable representing absence or presence of the bat species, labeled by 0 and 1 respectively. Let ℐ be the cells index set of study area, and let 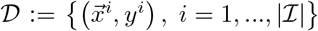 be the dataset, where let 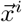 be to the predictor vector labeled by *y*^*i*^ ∈ {0, 1} and, *y*^*i*^ = 1 if it is a presence observation and *y*^*i*^ = 0 if it is an absence. When necessary, we will refer to *y*_*b*_ to the response variable corresponding to species *b* ∈ *ℬ*, where ℬ= {1, …, 25} (see Table S1). Notice that 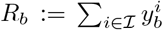 represents the occurrence/presence number of bat species *b* in our dataset, and therefore the Fig. 1 (top) shows its boxplot.

### 2.2 How RUSBoost works

RUSBoost method combines Random UnderSampling (RUS) and AdaBoostM1 algorithm. Specifically, RUS is a technique which randomly removes data points from the majority class, and AdaBoostM1 is a boosting technique used as an Ensemble Method (Freund et al., 1996) (see Fig. 2) which sequentially trains a next learner model on the mistakes of the previous learners. In mathematical terms, RUS extracts random subsets 𝒟_*l*_ (with replacement) of dataset 𝒟 of size 2 · *R* < |ℐ|, where *R* denotes the number of members in the class with the fewest members in the dataset. In our study, as the bat presence records are the minority class, then *R* will be set (for each model) to *R*_*b*_. Subsequently, AdaBoostM1 builds a model (or learner), denoted ℳ_*l*_, using 𝒟_*l*_ as sub-dataset (or training set), for *l* = 1, …, *L*, with *L* denoting the number of learners/sets. Later, classifiers ℳ_*l*_ are used in parallel, each one offering an opinion 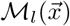 as to which class the example should be labeled with, to predict the most likely class for an unobserved vector 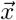. A “master classifier” (denoted by ℳ, see Fig. 2) collects this information and then chooses the label *y* that has received more votes as follows:

**Figure 2:**
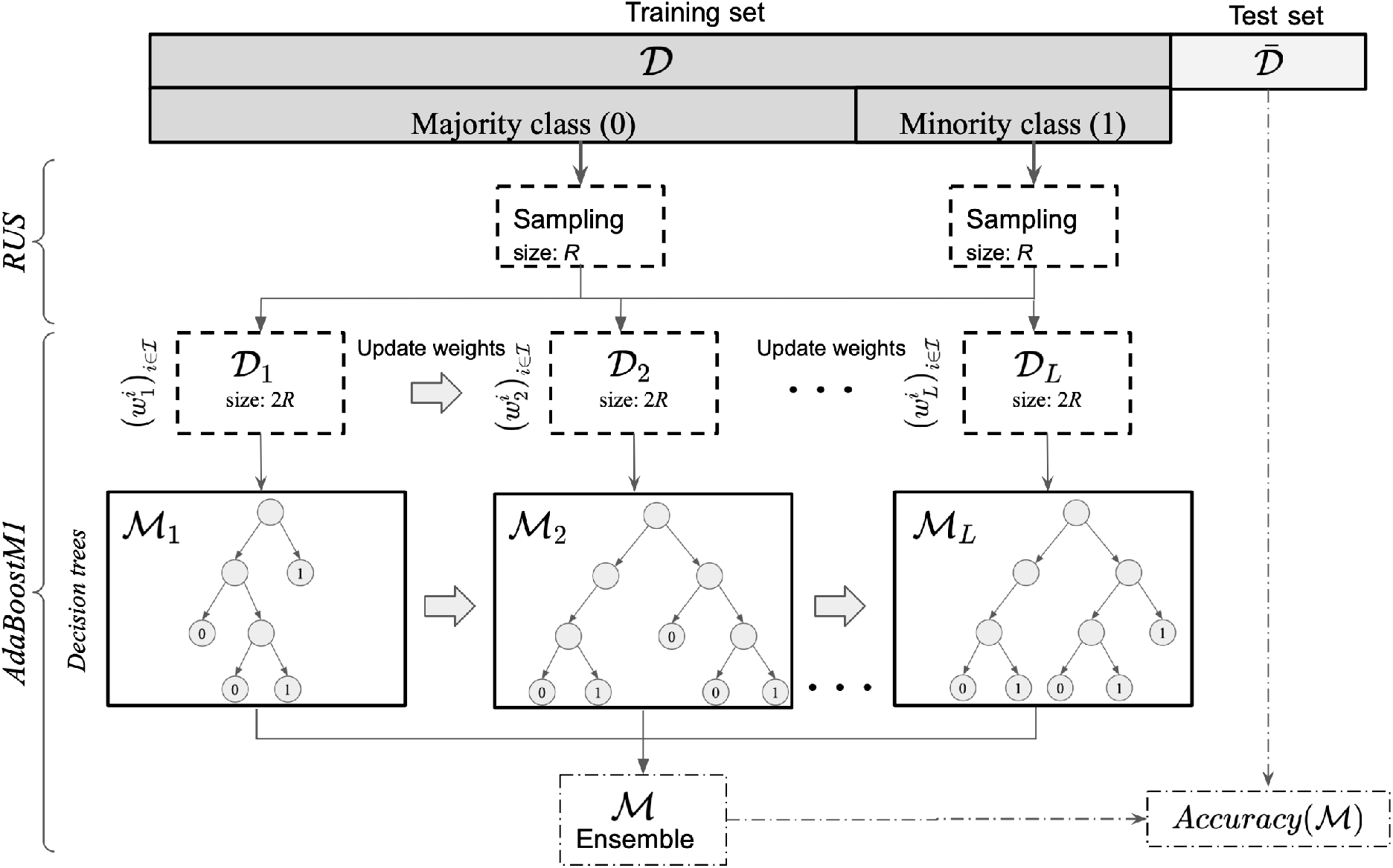
RUSBoost scheme: reading from top to bottom, the first *L* subsets of data are created from training sample 𝒟 using RUS procedure. Then, each subset data 𝒟_*l*_ is used to train their decision trees ℳ_*l*_ into the AdaBoostM1; and in this way, a set of different models are obtained sequentially. Finally, (ℳ_*l*_)_*l*=1:*L*_ models are combined into a “stronger ensemble” ℳ. The output response of ℳ is calculated by the Eq. (1) (right side). Using the data set 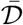 and Eq. (4), the accuracy level of ℳ is calculated.

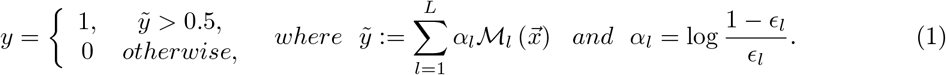

where the summation Eq. 1 can be seen as a weighted voting scheme and *ϵ*_*l*_ denotes the weighted classification error:

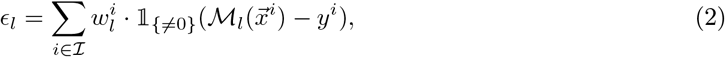

and 𝟙_{≠0}_(·) is the indicator function. In our implementation, we normalize the *α*_*l*_-parameters, so that 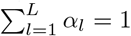, and we denoted 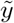 by 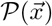, the *probability of bat occurrence* for the variable vector 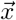.

In Fig. 2, for *i* ∈ ℐ, the weight 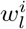 represents the probability that the observation pair 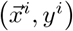 is drawn to train the *l*-th learner model. AdaBoostM1 increases weights for observations misclassified by learner ℳ_*l*_ and reduces (or keep them) weights for observations correctly classified. Then, next learner *l* + 1 is trained on the data with updated weights 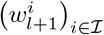. Models with *ϵ*_*l*_ ≥ 1*/*2 are discarded, and therefore the final modelℳ is only assembled with models ℳ_*l*_ that meet *ϵ*_*l*_ < 1*/*2, so that (1 − *ϵ*_*l*_)*/ ϵ*_*l*_ *>* 1, for all *l* = 1, …, *L*. From this, the procedure for updating the weights is as follows:

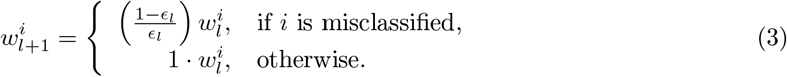

#### 2.2.1 Accuracy Assessment

After training the model, a validation accuracy score estimates the performance of the model on an independent data set 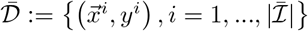. This procedure is known as *holdout validation*. It is worth mentioning that the full data set is such that 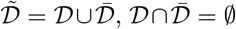; also that it is common to set 𝒟 to 75% of the entire dataset and 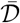 to 25%. Now, let ℳ be the ensemble model built following the procedure described in the previous section, using the training set 𝒟. Then, the accuracy of the method is defined by:

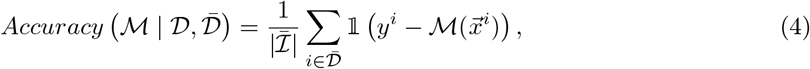

*Accuracy*, 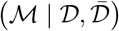 represents the percentage of successes obtained by the model on the validation dataset 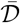. When the datasets of observations are divided into *k* disjoint subsamples (or folds), chosen randomly but with roughly equal size; then is taken a group as a hold out or test dataset 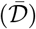 and the remaining groups as a training dataset (𝒟), this procedure is known as *K-Fold Cross-Validation*. In our study, we adopted the latter procedure (with *K* = 5)to validate and protect our model from overfitting and we estimate the average classification error as:

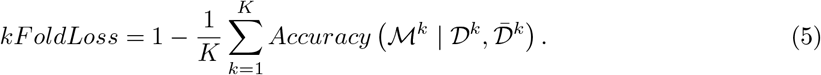

In addition, *specificity* and *sensitivity* are employed to appraise the classification capability computed in the following manner:

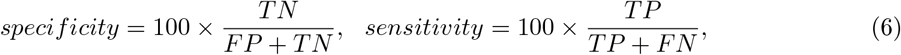

where TP (true positive) and TN (true negative) are the number of samples that are correctly classified as positive (presence class) and negative (absence class) observations in the cross validation process, respectively. FP (false positive) and FN (false negative) are the number of samples that are misclassified. Therefore, *sensitivity* is the percentage of positive (presence class) observations that are correctly classified whereas *specificity* is the percentage of negative (absence class) observations that are correctly identified. In addition, an area under the curve (AUC) of a receiver operating characteristic (ROC) curve was used for evaluating the averaged performances of classifiers (Breiman et al., 2001), which shows the true positive (TP) rate versus the false positive (FP) rate attained by the model for different thresholds of the classifier output. This way, we can monitor if the model is able to extract and learn the characteristics that distinguish bat presence from absences ones. The ROC chart for a good classifier tend to rise sharply at the origin and then level off near the maximum value of 1.0. Finally, we consider Area Under the ROC Curve (AUC), this metric is generally considered to be an important index to quantitatively assess the overall accuracy of the classifier’s performance. An AUC value near 0.5 means that the predictive ability of the model is completely random and a value of 1.0 represents a perfect prediction without misclassification. The closer the AUC value is to 1, the better the performance of the bat binary occurrence model.

#### 2.2.2 Hyper-parameter optimization

The performance of machine learning algorithms, broadly depend on parameters (named hyper-parameters) that are calibrated by the user’s expertise (which is difficult and time-consuming), recommendations from the algorithm package, or by more sophisticated methods that find the “best parameters” which improve some metric performance of the method (Yang and Shami, 2020). The hyper-parameters are internal parameters of any classifier or regression model and has a direct impact on the model’s performance. In this study, we used a Bayesian Optimization Method (BOM) (Eggensperger et al., 2013) to automate hyper-parameter settings used in the RUSBoost method (practical examples in Snoek et al. (2012)). In particular, we optimized the “Maximum Number of Splits (*MNS*)”, the “Number of Learners (*NL*)”, and the “Learning Rate (*LR*)” hyper-parameters. Higher *MNS* values lead to wider and more detailed decision trees as the number of leaves is increased, being able to identify more accurate split thresholds for the features. However, larger values may cause overfitting of the model to the training data in order to maximize its accuracy. Increasing the number of learners *NL* leads to a more time-consuming but potentially more accurate training of the model as the number of independent models learning from the data is increased. This allows the model to explore alternative classification schemes and focus on those difficult cases to adjust the weights of incorrectly classified instances (see *α*_*l*_ in the Eq. (1)). However, increasing the number of learners does not guarantee better performance (a priori). A natural trade-off between *NL* and *LR* arises as smaller learning rates lead to “slow down” the learning in the model, decreasing the contribution of each classifier. This plays a crucial role to avoid overfitting to the training data (a common drawback of decision tree based models), obtaining a more general model. On the other hand, larger *LR* values lead to faster convergence, potentially with the same or better performance than smaller rates. Therefore, *LR* acts as a weighting factor for how quickly the model learns from the classifiers, existing an inherent trade-off between these two parameters. Given the highly combinatorial nature of the problem, we require an efficient method to explore and evaluate promising combinations of these crucial hyper-parameters in practical running times.

Broadly, BOM attempts to minimize (or maximize) a scalar objective function 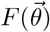 for 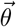 in a bounded domain Θ where lies of hyperparameters. No additional assumptions are made about *F*, and then *F* can be seen as a black-box function. In our study, the function 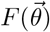 represents the cross-validation loss 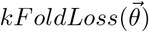 (see Eq. (5)) of the model (we set *K* = 5) and variables 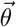 its hyper-parameters, and therefore, minimizing kFoldLoss function implies improving the predictive power of the model of both classes. The components/variables of 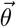 can be continuous reals, integers, or categorical. In our implementation let *θ*_*MNS*_ and *θ*_*NL*_ be the *MNS* and *NL* integer bounded variables; and *θ*_*LR*_ a *LR* real continuous bound variable. In particular, we arbitrarily assume that 10 ≤ *θ*_*MNS*_ ≤ 500, 1 ≤ *θ*_*NL*_ ≤ | 𝒟| − 1 and 10^−3^ ≤ *θ*_*RL*_ ≤ 1.0. Thus, Θ = [10, 500] × [1, | 𝒟| − 1] × [10^−3^, 1] ⊂ ℤ_+_ × ℤ_+_ × ℝ_+_. Therefore, the optimization problem solved is:

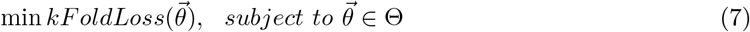

and whose optimal values we will denote by 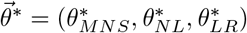.

### 2.3 Exploitation of RUSBoost for interpretability and decision making

A critical aspect of ML models is their interpretability (Castelvecchi, 2016). Despite the development of algorithms with remarkable performance, researchers and ecologists often fail to understand the logic behind the outputs; instead, they use them as an efficient black box. The previous sections are within this perspective, but we still need to understand what the role of variables is in predicting the probability of presence; or in the presence/absence classification process, and this is what we analyze in this section.

#### 2.3.1 Variable importance measure

There are several ways to determine the relative importance of variables and it depends mainly on how the method works. Variable importance in ensemble methods is derived from the feature importance provided by its weak learner (or base classifier) (see ℳ_*l*_ in the Fig. 2). In our approach we used a decision tree as a base classifier and then the AdaBoost feature importance was determined by the average feature importance provided by each decision tree.

“Predictor Importance (PI)” (Breiman, 1996) is a measure that assigns a score to each variable based on its contribution to the model learning process. With PI, candidate predictor variables can be compared according to their impact in predicting the response, or even their causal effect (Strobl et al., 2008). Our modeling approach allowed us to maintain all variables considered that could interact with bat occurrence. However, these values should not be understood as the estimated parameters as in the Linear Regression Models (LRMs), which act as weights in the predictions. For example, the interpretation of a weight in the linear regression model to the increase of a numerical feature by one unit changes the estimated outcome by its weight (in the case a numeric feature), which is not the interpretation in RUSBoost or others non-parametric methods. Also, variable importance scores only allows positive values, which does not occur in LRMs methods. The next section presents how to analyze the effect of a variable with respect to the response predicted by the model.

#### 2.3.2 Effect of a variable on presence probability

We used Partial Dependence Plots (PDPs) (Elith et al., 2008; Friedman, 2001) to analyze relationships between explanatory variables and bat occurrence. PDP shows, in particular, the partial contribution of a given variable considering all other variables at their measured values, presented visually by line plot (2D graphic); or via a 3D surface. For example, if we separate the attribute vector 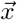 in a subset of attributes 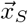 and 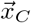, where the latter vector represents the complement over 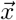, we get 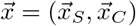 and the expectation of predicted answers regarding 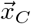 is naturally given by:

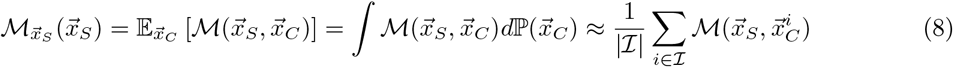

where 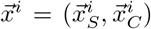 is the *i*-th observation. The last sum in Eq. (8) represents an approximation, assuming that each observation is equally likely. The PDP is defined as the graph of the function 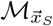, which we will denote by 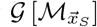. A PDP can show whether the relationship between the bat occurrence and a specific set of variables is linear, monotonic or more complex, and it can provide an important interpretability of how the variable to be evaluated is influencing (in our study) the probability of the species presence.

The summation in the Eq. (8) assumes the existence of the data points 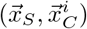, and then when the variables are correlated, such artificial samples may have a very low probability measure, therefore, before building our models, in the database of our case study, we eliminate highly correlated variables (*>* 0.5).

## 3 Results

### 3.1 Hyper-parameter tuning and modeling

We built models for each bat species (|ℬ|= 25) and obtained the values of the hyperparameters after the optimization process (see Table S1 of Supplementary Material for more details). After cross validation procedure, the averages values for AUC, specificity, sensitivity, and overall accuracy were 0.8 ± 0.1, 79.6 ± 4.9%, 78.8 ± 5.0%, and 78.8 ± 5.0%, respectively (see Fig. 3 and Table S1), and therefore, models predicted species presence better than random (AUC values *>* 0.5). The average values obtained for *MSN, NL* and *LR* were 2402.7 ± 2451.7, 446.24 ± 102.92 and 0.1 ± 0.2, respectively (see Table S1).

**Figure 3:**
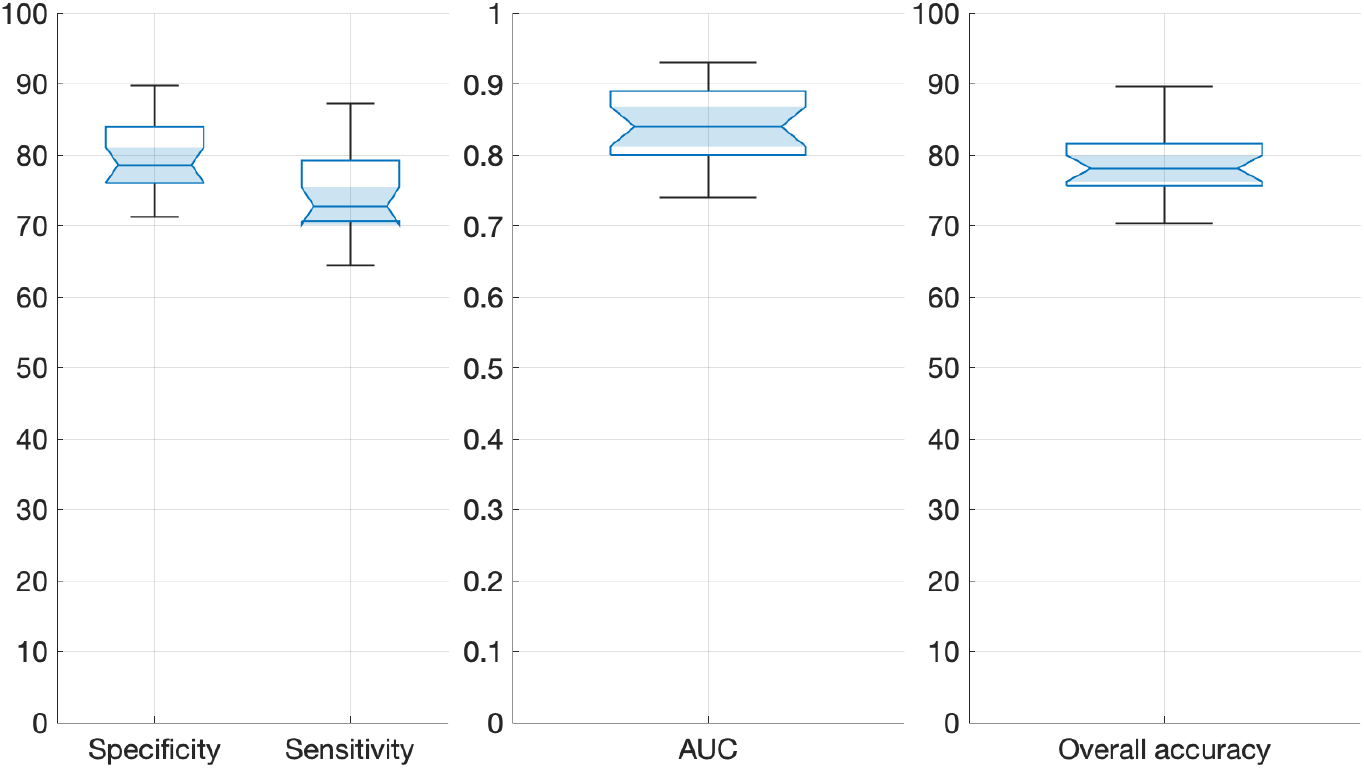
Summary of performance models, where the left and right chart are expressed in percentage and the center in a range from 0 to 1.

We analyzed the effect of the hyperparameters optimization procedure using the bat species with most and least presence. Therefore, we used the distribution of *P. pipistrellus* with a total presence of 37.2% (2297 records) and *P. nathusii* with a presence of 0.5% (33 records) (Fig. 5). In this particular case, the bayesian method iteratively explored the values until converging to a solution that minimized the the *kFoldLoss* described in the Eq. (5). Fig. 4 shows the best hyper-parameter values obtained when solving the optimization problem defined in the Eq. (7) for the species *P. pipistrellus*: 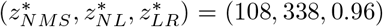; and *P. nathusii* : 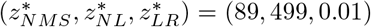. These values were reached after iteration 23 for the model corresponding to Ppip (Fig. 4 - left side) and 13 for Pnat species (Fig. 4 - right side).

**Figure 4:**
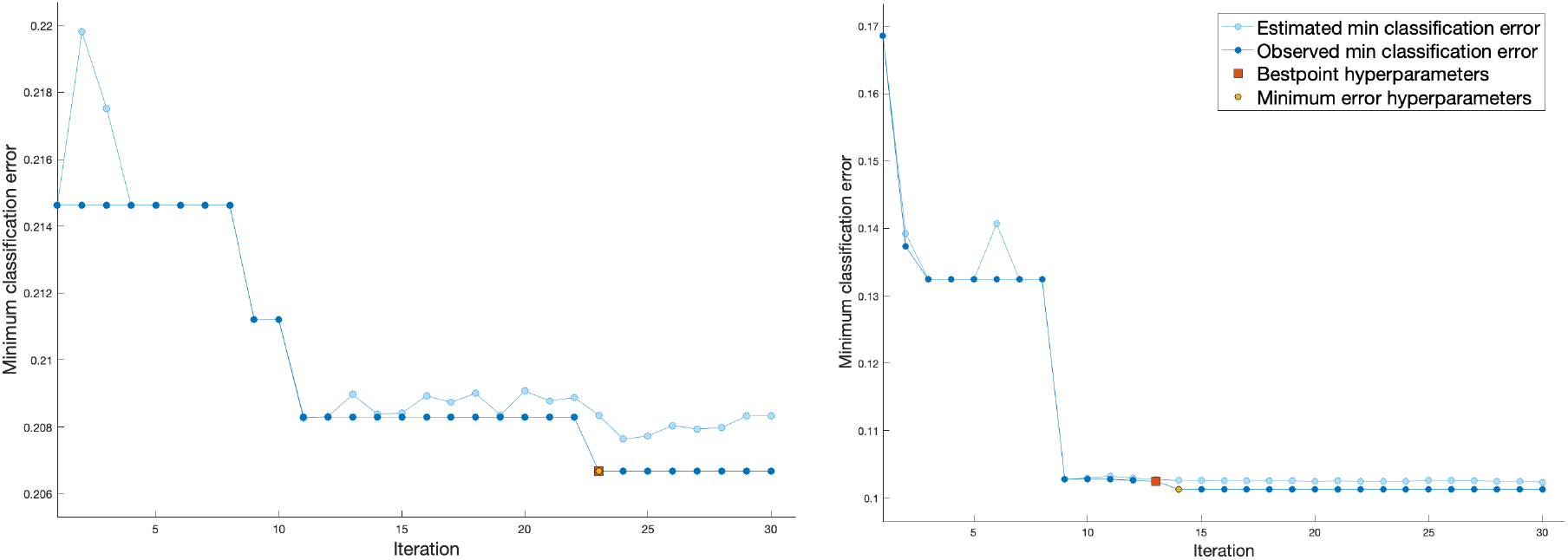
Minimum classification error (MCE) evolution for Ppip (left chart) and Pnat (right chart). The MCE is updated as the optimization algorithm runs. Each light blue dot represents an estimate of the MCE value for a combination of hyper-parameter values 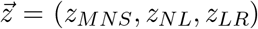 tested until the current iteration. Similarly, dark blue dots indicate the observed minimum MCE. The red square and orange dot (the two marks coincide on the left graphic) indicate the best hyperparameters combination and those that produce the minimum observed MCE, respectively.

**Figure 5:**
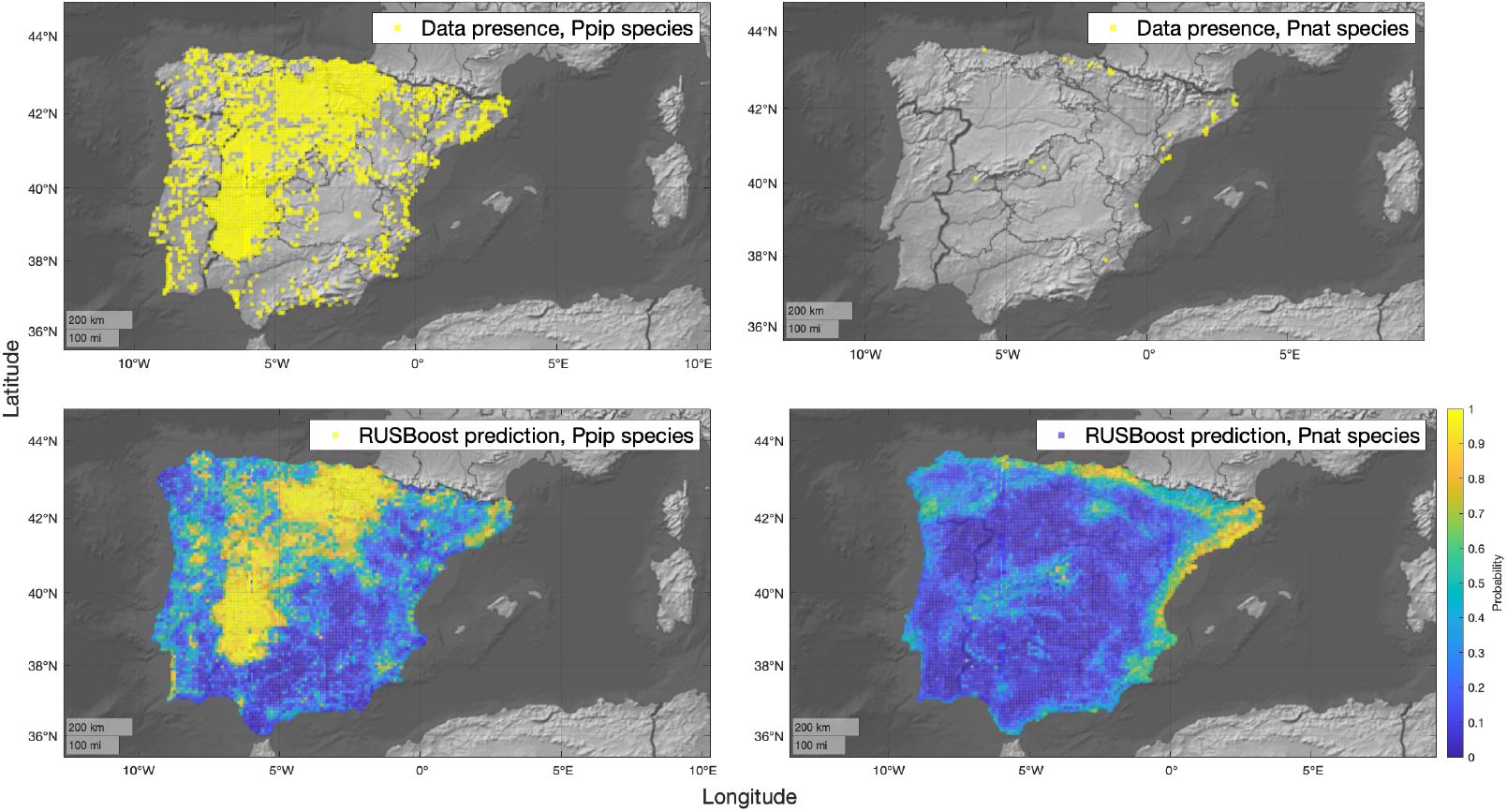
RUSBoost estimations for Ppip and Pnat species

The minimum classification error (MCE) decreases with the number of iterations, until it stabilizes at an “optimal” value (Fig. 4).

Once the hyperparameters 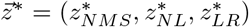 have been established for each model/species, the ℳ^*Ppip*^ and ℳ^*Pnat*^ models are built and then used to calculate the presence probability in each cell of our study area (a total of 6,169 cells within Iberian Peninsula); Fig. 5 shows the result of this procedure. The maps located in the upper show the presence records of each species and the maps located in the lower the probabilities predicted by each model. Visually, we can see that presences and high values are properly correlated. In our Supplementary Material we present the probability maps for the rest.

We observed performance metrics from Table S1, that after cross validation procedure, the averages values for AUC, specificity, sensitivity, and overall accuracy were 0.93, 89.7%, 78.8%, and 89.6%, for Pnat species respectively; while that for Ppip were 0.86, 84.0%, 71.0%, and 79.1%, respectively. In both cases, good values were obtained for all the metrics, however, the values were not proportional to the number of presence records. Broadly, we did not find a significant correlation between the number of species records with the performance measures of RUSBoost method.

### 3.2 Importance of the variables

This section highlights the variable importance estimations in the construction of the models. In Fig. S1 of our Supplementary material the values obtained for each model/species are shown, of the thirty-six variables considered in our study, and for greater clarity, we summarized the results in the Fig. 6 and 7. Specifically, the first figure shows the variables corresponding to land cover (left chart) and weathers (right chart); and the second corresponds to topography and human activity. According to the figures, the land covers with the highest value of importance in the prediction of most of the models were the variables *Forest, Shurbland* and *Rainfallcrops*, whose medians exceed 3%. These variables are closely followed (below 3%) by *Pasture, Mixedcrops, Abandoned, Irrigatedcrops* and *Urban* covers. The remaining variables such as *Mines, Roads, River, Wetland, SandRock, Agroforestry* and *Improductive* are below the 2% importance value according to our results.

**Figure 6:**
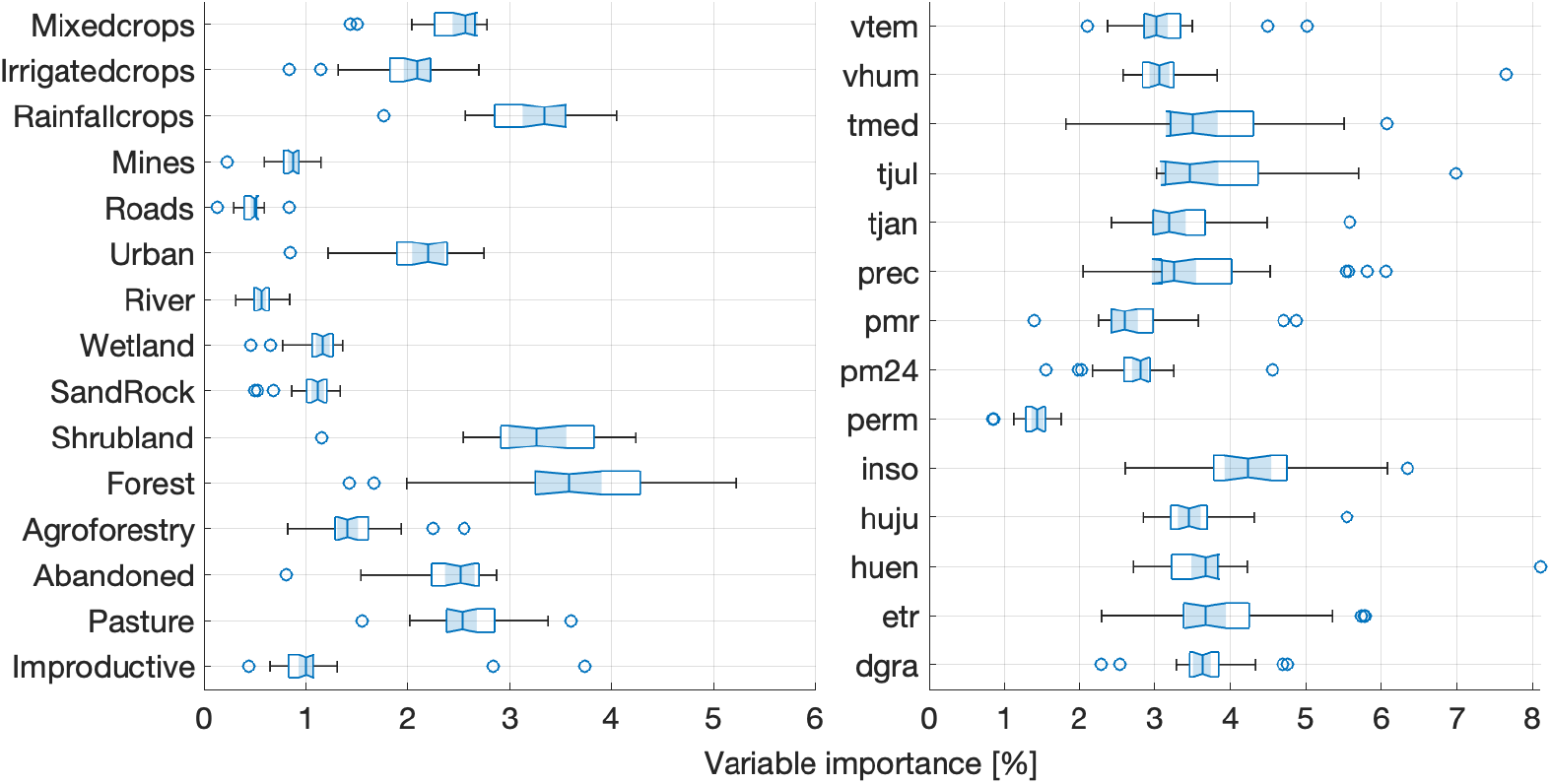
Importance of variables of Land cover and Weather drivers.

**Figure 7:**
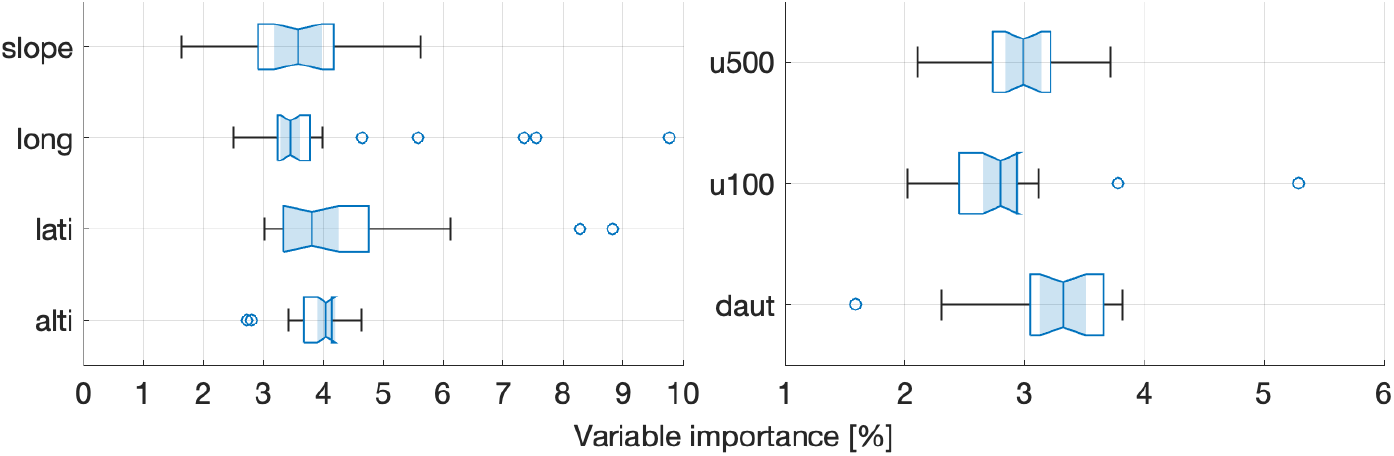
Importance of variables of Topography and Human Activity drivers.

Regarding to weather drivers, most are in the range of 3% to 5%, except for the variables *pmr, pm*24 and *perm* that represent the relative maximum precipitation; 24-hour maximum rainfall; and soil permeability respectively. Only the variable *inso* exceeds 4% of importance (its median), and then the variables of relative humidity in January (*huen*), average annual evapotranspiration (*etr*), average annual temperature (*tmed*), average July temperature (*tjul*), and so forth.

Finally, in Fig. 7 presents the values obtained for topography (left chart) and human activity (right chart) drivers. All topographic variables are in the range of 3 to 5 percent, although their medians are concentrated below 4%. On the other hand, in the human activity variables it can be appreciated that only the *daut* variable exceeds 3% while the other variables (*u*100 and *u*500) are below this value.

## 4 Discussion

### 4.1 Handling class imbalance

The ability of handling class imbalance is a necessary skill in our study, since the variability of record numbers for each bat on each trained the model is small, due to the low sighting frequency of each species (see Fig. 1 and Table S1). In our study, we have applied RUSBoost to a data of presence/absence of bats in a vast region and with very uneven and imbalanced records. However, the method, independently of the number of records, obtained an excellent performance (see Fig. 8), comparable to other similar studies (see e.g. Maher et al. (2014)).

**Figure 8:**
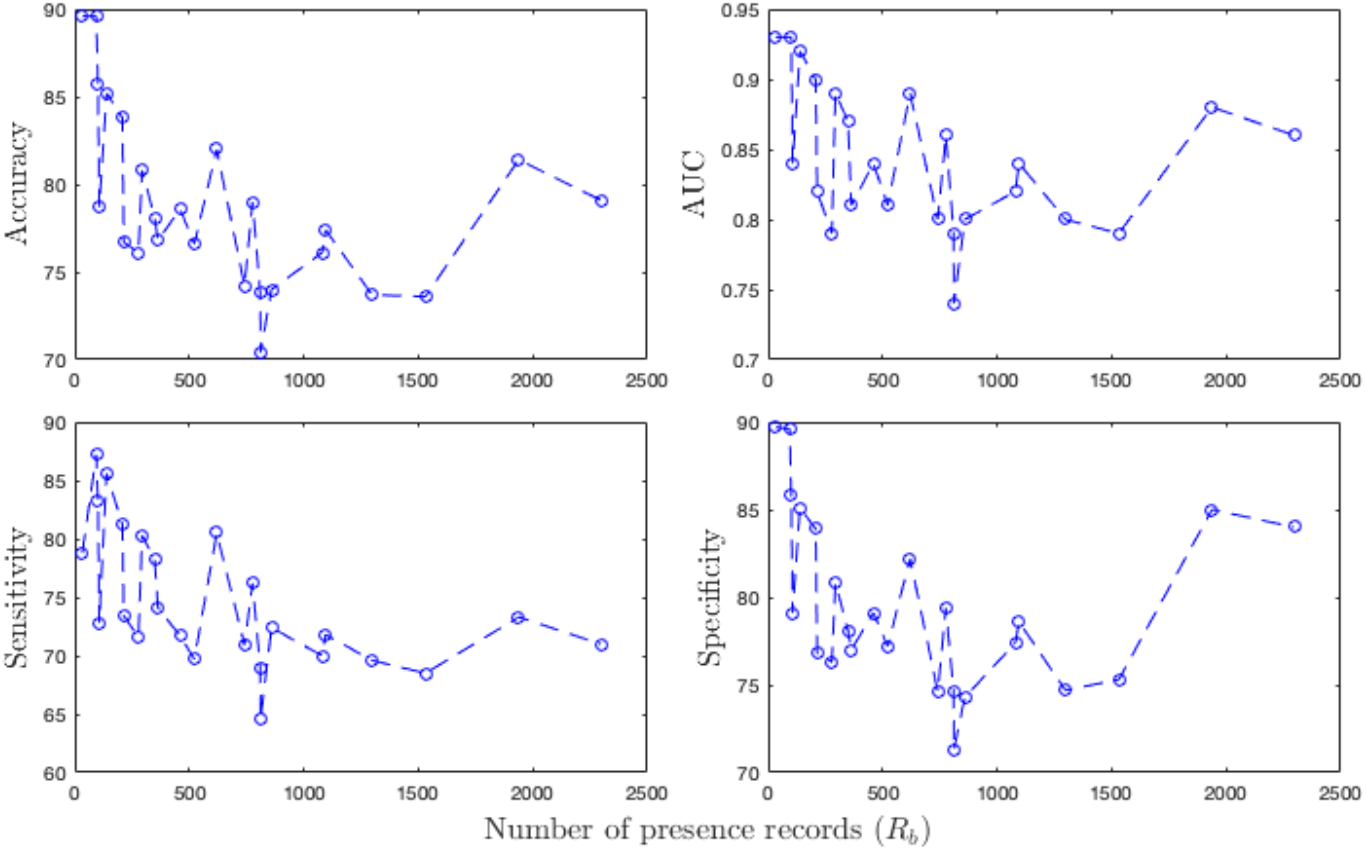
Number of presence records vs performance metrics.

Applying methods such as Random Forest or AdaBoost are not suitable when classes in categorical response variables are imbalanced (Chen, 2004) due to bootstrap over-representing the majority class, leading to under-prediction of the minority class, so the overall performance measures (AUC and Overall Accuracy) can be misleading. Currently, there are three ways to deal with imbalanced data: i) assigning a high cost to minority class misclassification (Chen, 2004); ii) reducing the sample of the majority class (Kubat et al., 1998); and iii) oversampling the minority class (Chawla et al., 2002). RUSBoost method relies on the second approach balancing the response variable that iteratively down-samples the majority class by randomly drawing 2 · *R* (where *R* is the size of minority class, see Fig. 2) from the majority class and building decision tree models iteratively using different random subsets while holding the sample-size of the minority-class constant (Evans and Cushman, 2009).

The RUSBoost method has been successfully applied in different contexts: to determine impervious surface areas (Kesikoglu et al., 2016); to identify dolphins species from characteristics extracted from photos (Maglietta et al., 2018); and was recently used to detect electrical losses in power systems Adil et al. (2020), which is compared with state-of-the-art methods showing the superiority for handling imbalanced data, parameter optimization and overfitting. Also, recently RUSBoost was used to assess risks of fire (or fire occurrence probability) for territorial planning (Carrasco et al., 2021), but the dataset was slightly imbalanced and contained only a type of “species” (fire and non-fire). However, to our knowledge in few references we have been able to find its use in species distribution models in a few references. A review of dedicated classification methods when the database is imbalanced can be found in Haixiang et al. (2017).

### 4.2 Hyper-parameter optimization

Every machine learning algorithm or software has hyper-parameters, and a desirable task is to automate them in order to optimize model performance. However, this is not an easy task, and the complexity of this process will mainly depend on the nature of the variables/hyper-parameters and the objective function to be optimized (e.g. performance measure), where the latter is usually a black-box function. Perhaps the most common algorithms to perform this task are: Grid Search (Bergstra et al., 2011), Random Search (Bergstra and Bengio, 2012), Genetic Algorithm (Lessmann et al., 2005) and Bayesian Optimization method (BOM) (Eggensperger et al., 2013). Contrasting with traditional optimization methods such as gradient descent based, for example: NEWUOA (Powell, 2006), BOBYQA (Powell, 2009), DFO (Conn and Toint, 1996), and so forth; Many of these algorithms also have the capacity to effectively identify discrete, categorical, and conditional hyper-parameters.

In algorithmic and very general terms, Grid Search (GS) is based on the concept of defining a fixed hyper-parameter search space and then detecting the hyper-parameter combinations in it, in order to find the best performing hyper-parameter combination. On the other hand, Random search (RS) randomly selects combinations of hyper-parameters in the search space (which may not be fixed), given limited execution time and resources. In GS and RS, each hyper-parameter configuration is treated independently and therefore can easily be parallelized. Unlike GS and RS, BOM determines the next hyper-parameter value based on the previous results of tested hyper-parameter values, which avoids many unnecessary evaluations; thus, BOM can detect the optimal hyper-parameter combination within fewer iterations rather than GS and RS (Yang and Shami, 2020). A comprehensive review of algorithms and applications can be found in Yu and Zhu (2020).

Broadly, there is no criterion to evaluate if the solution (combination of hyper-parameters found) is the optimal solution of the optimization problem defined in the Eq. (7) because the *kFoldLoss* function (see Eq. (5)) is a black-box function and therefore we do not know about its structural mathematical properties (continuity, differentiability, etc). On the other hand, the number of iterations is predefined by the user, in our study in particular, a maximum number of 30 iterations were arbitrarily established, due to the computational time consumption (approximately one minute per iteration).

#### 4.2.1 Variable interaction on presence probability

In this section, we discuss the potential of our methodology to analyze the probability sensitivity of bat occurrence, with respect to the variation of a variable through an illustrative example (see Section 2.3.2). For them, we built the functions 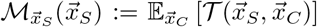 for *S* representing individual variables, i.e. |*S*| = 1 (see Eq. (8)). Strictly speaking, we could have carried out this analysis for each of the variables and for each of the species considered in our study, producing a total of 36 × 25 = 900 PDPs. However, for practical and expository purposes, we only include the analysis of the effect of the variable *tjul* on the probability of the presence of the species *Barbastella barbastellus* (Bbar).

Because RUSBoost algorithm involves Random Undersampling of data for building the ensemble model detailed rule sets can vary slightly between different model runs, and therefore it is advisable to run multiple bootstrap iterations of algorithm to achieve stable and repeatable results (Murphy et al., 2010). The influence of the temperature variable on the predicted presence probability is visualized in Fig. 9. The plot shows a slight decrease in probability from 14 to 21 Celsius’ degrees, and a steep decrease from 21 to 26 degrees. Above 26 degrees the probability stabilizes. The figure also shows that the probability values remain within an acceptable error (see standard deviation in the chart), under different replications for the construction of the model, and therefore the mean values (blue-line) can be taken as a reliable estimate.

**Figure 9:**
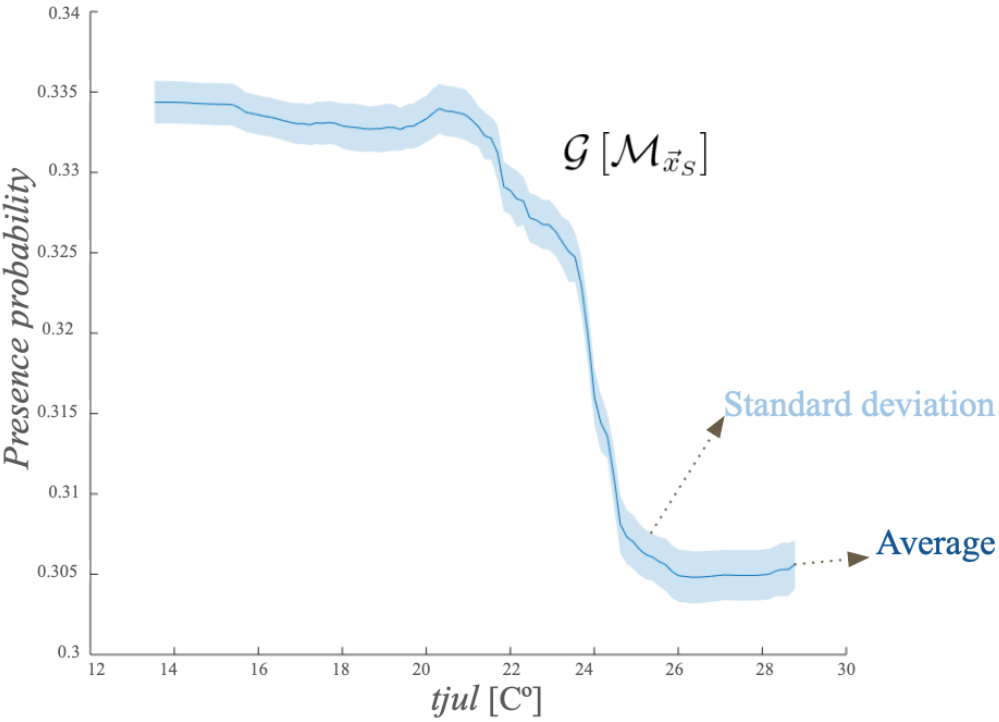
Partial dependence plot for *tjul* variable and *Bbar* species.

#### 4.2.2 Bayesian interpretation

From Eq. (1), weighted voting schemes can be seen as approximations under a Bayesian approach. In fact, let 𝒴 be the random variable which represents the choice of the presence/absence class, and informally we denote ℙ (ℳ_*l*_) = *α*_*l*_ as the discrete probability distribution selection of decision trees in the RUSBoost procedure. Replacing 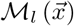 in the Eq. (1) with the class posterior probabilities denoted by 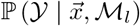 (or relative frequency of classes, before the tree leaf in the decision tree model ℳ_*l*_). From this, the Eq. (1) (right side) can be rewritten as follows:

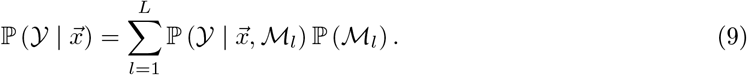

Eq. (9) helps understand the intuition behind the previously defined probability of bat occurrence (see Eq. (1)) as a probability that is conditional on the variables of a observed point 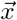, and it is obtained as a weighted summation of the class posterior and decision trees selection probabilities.

## 5 Conclusions

The RUSBoost method assessed in this study yielded similar performance values (AUC, sensitivity, specificity and overall accuracy) to common techniques and, thus, expand modelling options for ecologists. We found that the performance of the method is invariant to the number of presence records (see Fig. 8) and therefore it can be applied as a method based on presence-only data, without losing the underlying abilities of the presence/absence methods, for example, enabling analyses of biases and prevalence (Phillips et al., 2009). It is important to emphasize that the method is suitable when the absence data is reliable. When this is not the case or the species not easily detected, it is preferable to use presence-only techniques such as MaxEnt (Phillips et al., 2006), GARP (Genetic Algorithm for Rule-set Production)(Stockwell, 1999) or SVMs (Support Vector Machines) (Guo et al., 2005). The method showed broadly a good balance between sensitivity and specificity, a very important aspect for rare and threatened species and conservation decisions (Guisan et al., 2013).

We also show that the method is sensitive to its hyper-parameters. However, they can be optimized with the Bayesian Optimization Method in a reasonable time. In the case of the model built for the Pnat species, the overall precision of the method increased from 83% to 90% in less than 15 iterations (approximately 15 minutes on our computer), which shows the advantages of combining the RUSBoost with hyper-parameter optimization.

## Authors’ contributions

J.C., F.L. and A.W. conceived the project; F.L. collected the data; J.C. and F.L. raised the scientific hypotheses and methodology; J.C. wrote the code; all authors contributed to writing the manuscript, with J.C. having the lead on the writing. F.L. and A.W. reviewed and edited the final manuscript.

## Acknowledgements

We thank Ángeles Haz for check and correct the English version of the manuscript. Jaime Carrasco acknowledges the support of the Agencia Nacional de Investigación y Desarrollo (ANID), Chile, through funding Postdoctoral Fondecyt project 3210311. Fulgencio Lisón was supported by Fondecyt iniciacion project N° 11180514, Andrés Weintraub thanks Fondecyt 1170381.

