## Supplementary Material for "RUSBoost: A suitable species distribution method for imbalanced records of presence and absence. A case study of twenty-five species of Iberian bats"

### Suplementary Material

#### Models performance

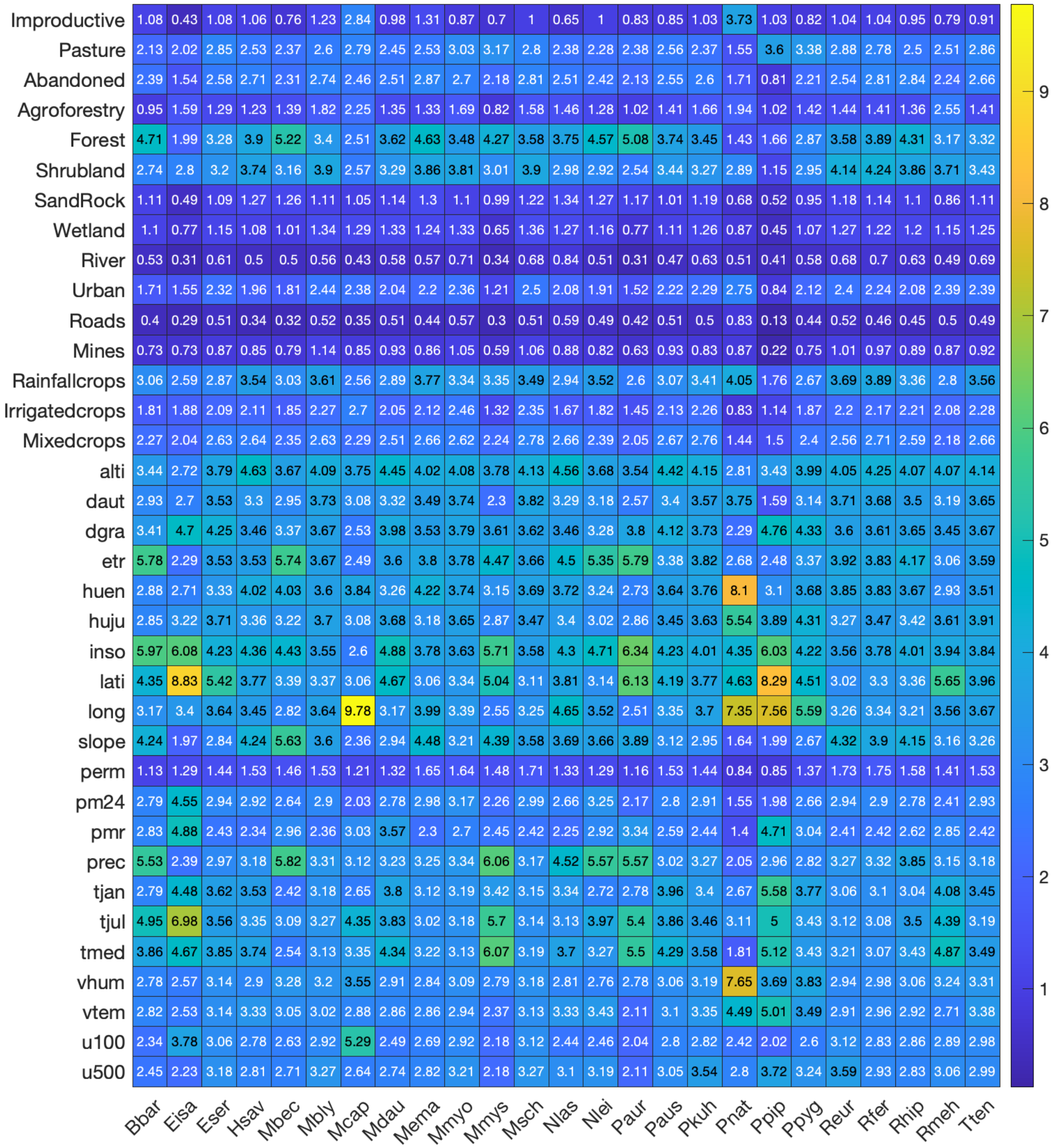


Figure S1: Importance of the variables obtained by RUSBoost models.

| **Species** | Acronisms ($b \in \mathcal{B}$) | **Records** | $\theta^*_{MNS}$ | $\theta^*_{NL}$ | $\theta^*_{LR}$ | **Specificity** | **Sensitivity** | **AUC** | **Accuracy** |
|---|---|---|---|---|---|---|---|---|---|
| Barbastella barbastellus | Bbar (1) | 349 | 1117 | 224 | 0.02 | 78.1 | 78.2 | 0.87 | 78.1 |
| Eptesicus isabellinus | Eisa (2) | 139 | 440 | 170 | 0.02 | 85.1 | 85.6 | 0.92 | 85.2 |
| Eptesicus serotinus | Eser (3) | 776 | 5828 | 255 | 0.02 | 79.4 | 76.2 | 0.86 | 79 |
| Hypsugo savii | Hsav (4) | 460 | 6052 | 497 | 0.02 | 79.1 | 71.7 | 0.84 | 78.6 |
| Myotis bechsteinii | Mbec (5) | 107 | 1656 | 500 | 0.02 | 79.1 | 72.8 | 0.84 | 78.7 |
| Myotis blythii | Mbly (6) | 278 | 1004 | 500 | 0.02 | 76.3 | 71.6 | 0.79 | 76.1 |
| Myotis capaccinii | Mcap (7) | 94 | 6127 | 500 | 0.02 | 89.6 | 872 | 0.93 | 89.6 |
| Myotis daubentonii | Mdau (8) | 859 | 4626 | 449 | 0.01 | 74.3 | 72.4 | 0.8 | 74 |
| Myotis emarginatus | Mema (9) | 363 | 373 | 500 | 0.02 | 77 | 74.1 | 0.81 | 76.8 |
| Myotis myotis | Mmyo (10) | 811 | 602 | 498 | 0.01 | 71.3 | 64.5 | 0.74 | 70.4 |
| Myotis mystacinus | Mmys (11) | 101 | 6056 | 480 | 0.02 | 85.8 | 83.2 | 0.93 | 85.7 |
| Miniopterus schreibersii | Msch (12) | 810 | 1309 | 500 | 0.02 | 74.6 | 68.9 | 0.79 | 73.8 |
| Nyctalus lasiopterus | Nlas (13) | 208 | 5832 | 500 | 0.02 | 83.9 | 81.2 | 0.9 | 83.8 |
| Nyctalus leisleri | Nlei (14) | 620 | 6107 | 491 | 0.01 | 82.2 | 80.5 | 0.89 | 82 |
| Plecotus austriacus | Paur (15) | 294 | 1259 | 498 | 0.01 | 80.8 | 80.3 | 0.89 | 80.8 |
| Plecotus auritus | Paus (16) | 741 | 375 | 496 | 0.01 | 74.6 | 70.9 | 0.8 | 74.2 |
| Pipistrellus kuhlii | Pkuh (17) | 1095 | 1103 | 497 | 0.01 | 78.6 | 71.8 | 0.84 | 77.4 |
| Pipistrellus nathusii | Pnat (18) | 33 | 89 | 499 | 0.01 | 89.7 | 78.8 | 0.93 | 89.6 |
| Pipistrellus pipistrellus | Ppip (19) | 2297 | 108 | 338 | 0.96 | 84.0 | 71.0 | 0.86 | 79.1 |
| Pipistrellus pygmaeus | Ppyg (20) | 1938 | 769 | 496 | 0.01 | 85 | 73.3 | 0.88 | 81.4 |
| Rhinolophus euryale | Reur (21) | 522 | 533 | 499 | 0.01 | 77.2 | 69.7 | 0.81 | 76.6 |
| Rhinolophus ferrumequinum | Rfer (22) | 1531 | 6109 | 500 | 0.01 | 75.3 | 68.5 | 0.79 | 73.6 |
| Rhinolophus hipposideros | Rhip (23) | 1298 | 452 | 499 | 0.01 | 74.7 | 69.6 | 0.8 | 73.7 |
| Rhinolophus mehelyi | Rmeh (24) | 214 | 1114 | 498 | 0.01 | 76.8 | 73.4 | 0.82 | 76.7 |
| Tadarida teniotis | Tten (25) | 1080 | 1027 | 272 | 0.02 | 77.4 | 70 | 0.82 | 76.1 |
| | $AVG$ : | 680.72 | 2402.7 | 446.24 | 0.1 | 79.6 | 74.6 | 0.8 | 78.8 |
| | $STD$ : | 594.08 | 2451.7 | 102.92 | 0.2 | 4.9 | 5.8 | 0.1 | 5.0 |

Table S1: Performance and hyper-parameter optimization of RUSBoost models.

### Maps of probability presence

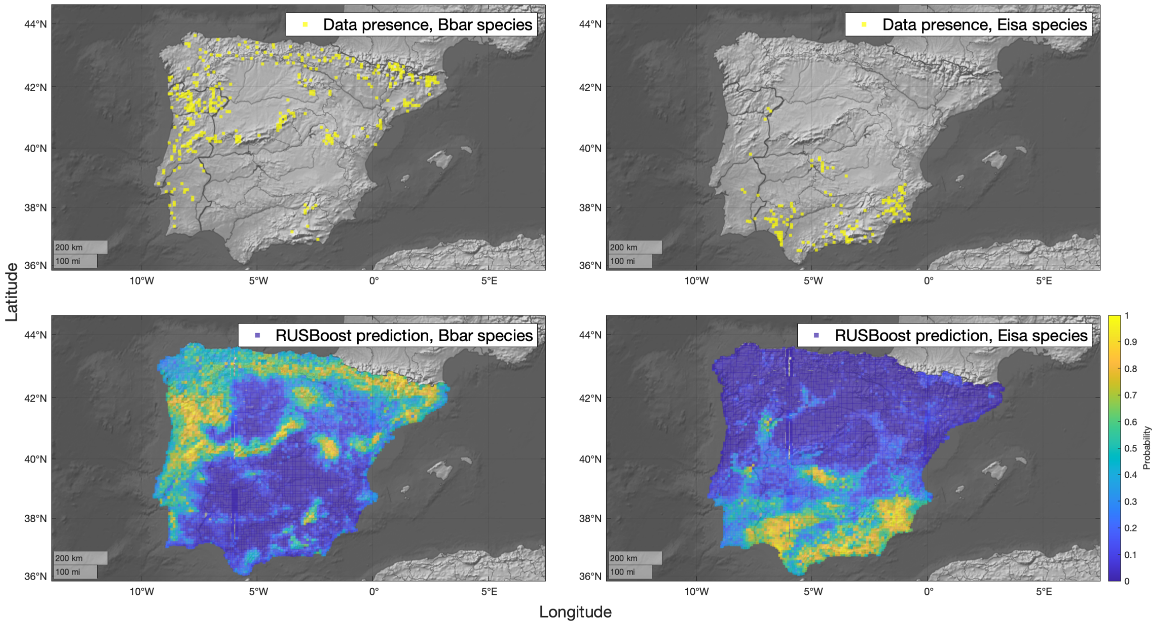


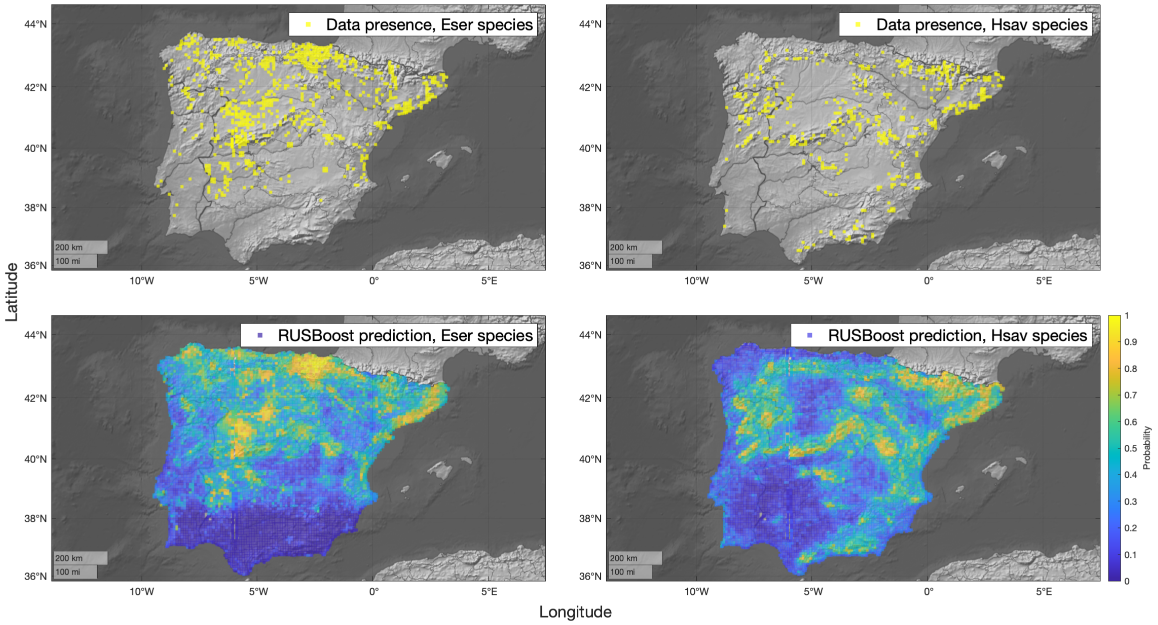

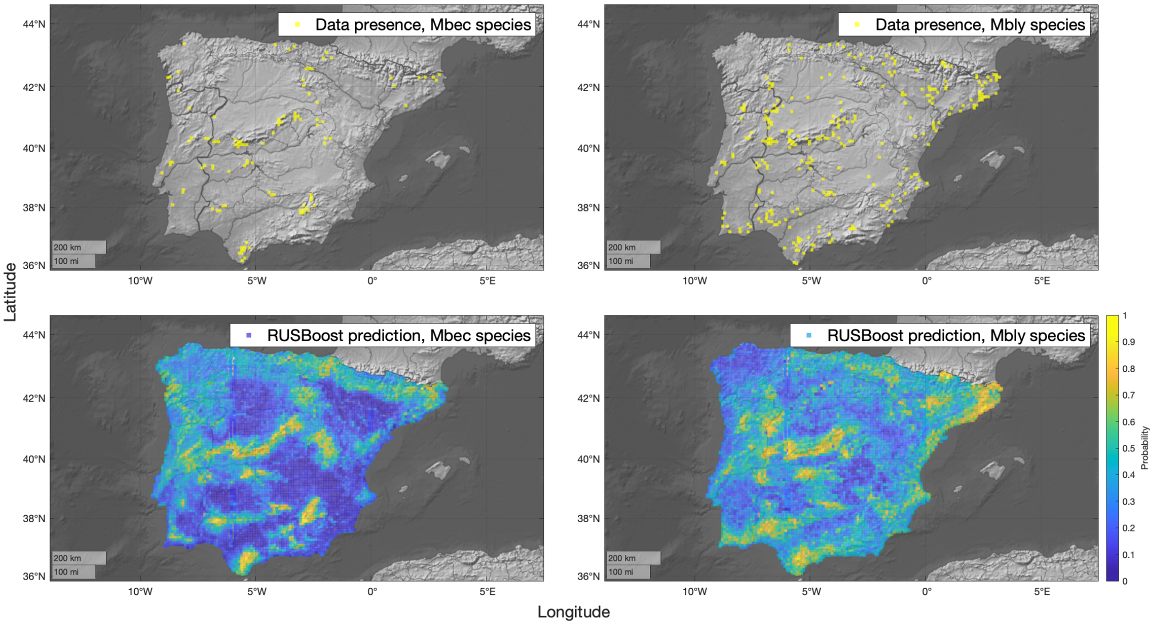
Data presence, Mbec species
Data presence, Mbly species
RUSBoost prediction, Mbec species
RUSBoost prediction, Mbly species
Latitude
Longitude
Probability
44°N
42°N
40°N
38°N
36°N
10°W
5°W
0°
5°E
200 km
100 mi
0
0.1
0.2
0.3
0.4
0.5
0.6
0.7
0.8
0.9
1

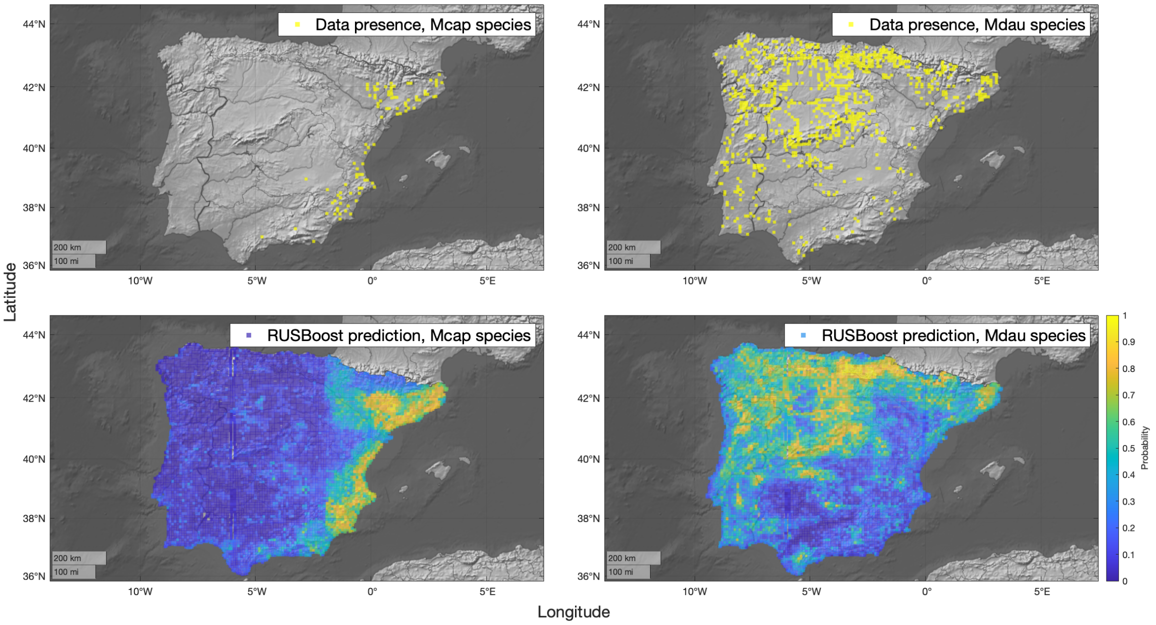
Data presence, Mcap species
Data presence, Mdau species
RUSBoost prediction, Mcap species
RUSBoost prediction, Mdau species
Latitude
Longitude
Probability
44°N
42°N
40°N
38°N
36°N
10°W
5°W
0°
5°E
200 km
100 mi
0
0.1
0.2
0.3
0.4
0.5
0.6
0.7
0.8
0.9
1

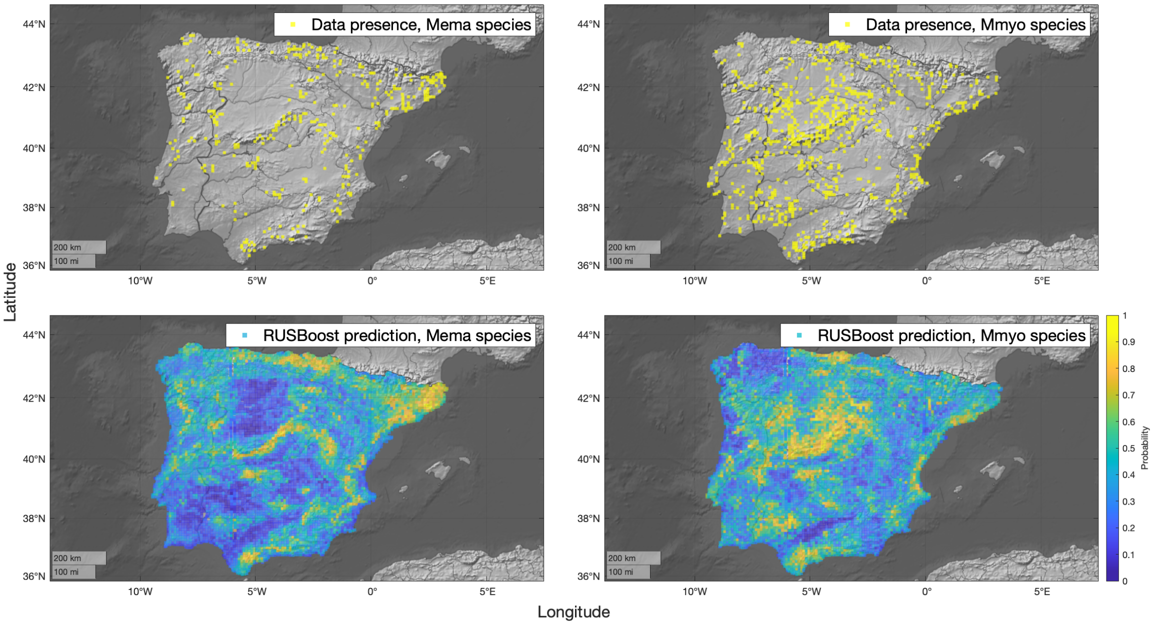
Data presence, Mema species
Data presence, Mmyo species
RUSBoost prediction, Mema species
RUSBoost prediction, Mmyo species
44°N
42°N
40°N
38°N
36°N
10°W
5°W
0°
5°E
200 km
100 mi
Latitude
Longitude
Probability
0
0.1
0.2
0.3
0.4
0.5
0.6
0.7
0.8
0.9
1

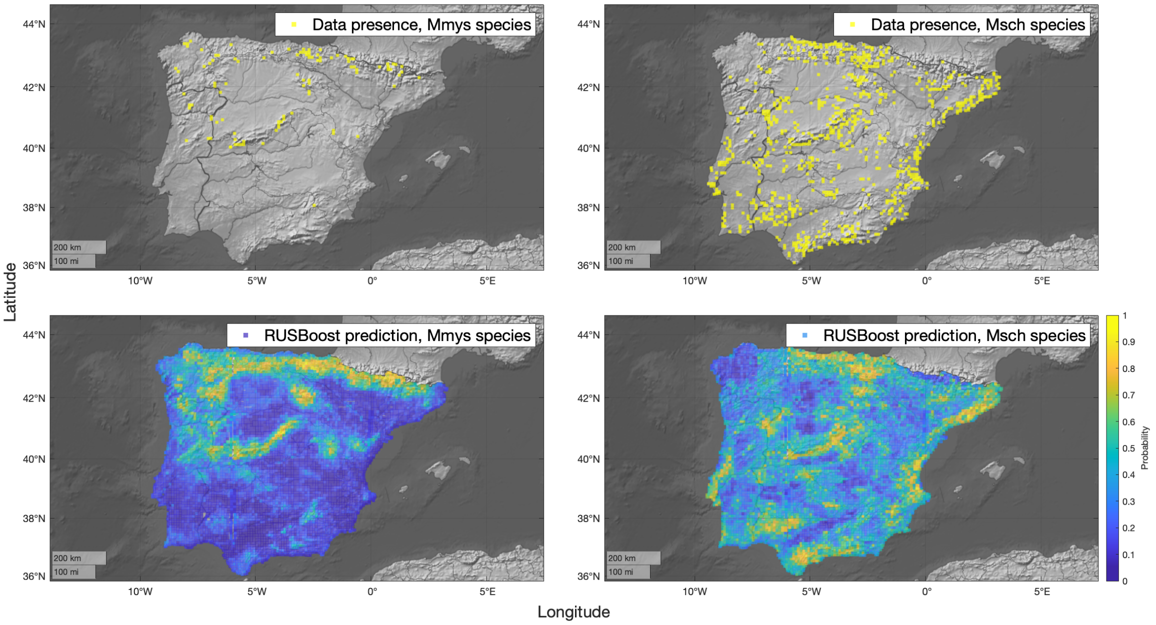
Data presence, Mmys species
Data presence, Msch species
RUSBoost prediction, Mmys species
RUSBoost prediction, Msch species
44°N
42°N
40°N
38°N
36°N
10°W
5°W
0°
5°E
200 km
100 mi
Latitude
Longitude
Probability
0
0.1
0.2
0.3
0.4
0.5
0.6
0.7
0.8
0.9
1

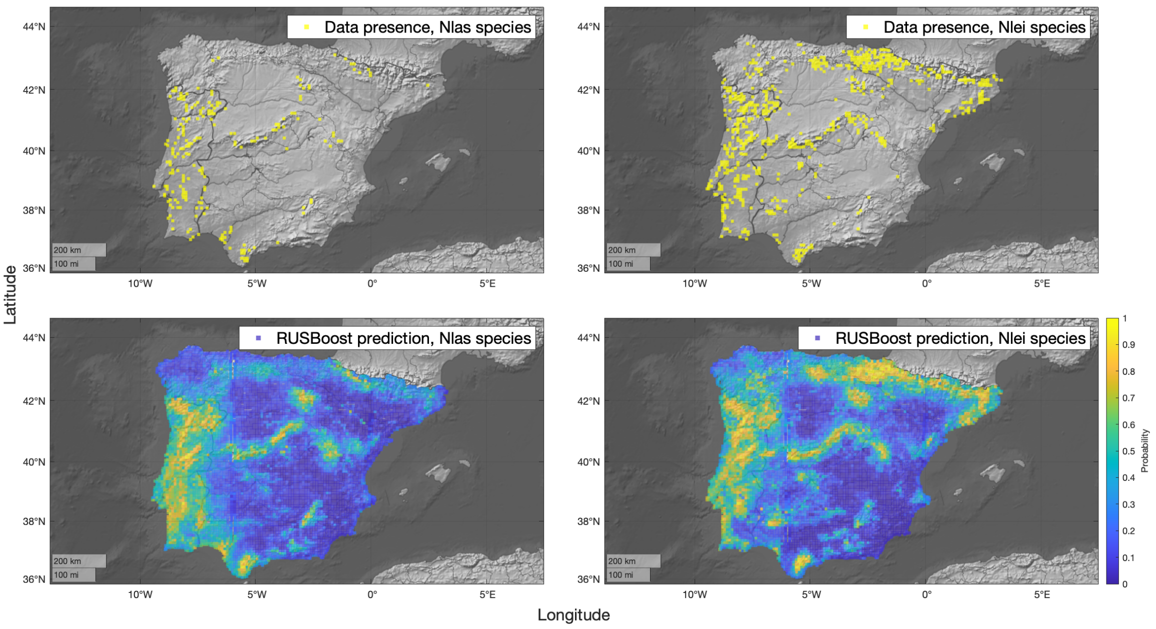
Data presence, Nlas species
Data presence, Nlei species
RUSBoost prediction, Nlas species
RUSBoost prediction, Nlei species
44°N
42°N
40°N
38°N
36°N
10°W
5°W
0°
5°E
200 km
100 mi
Latitude
Longitude
Probability
0
0.1
0.2
0.3
0.4
0.5
0.6
0.7
0.8
0.9
1

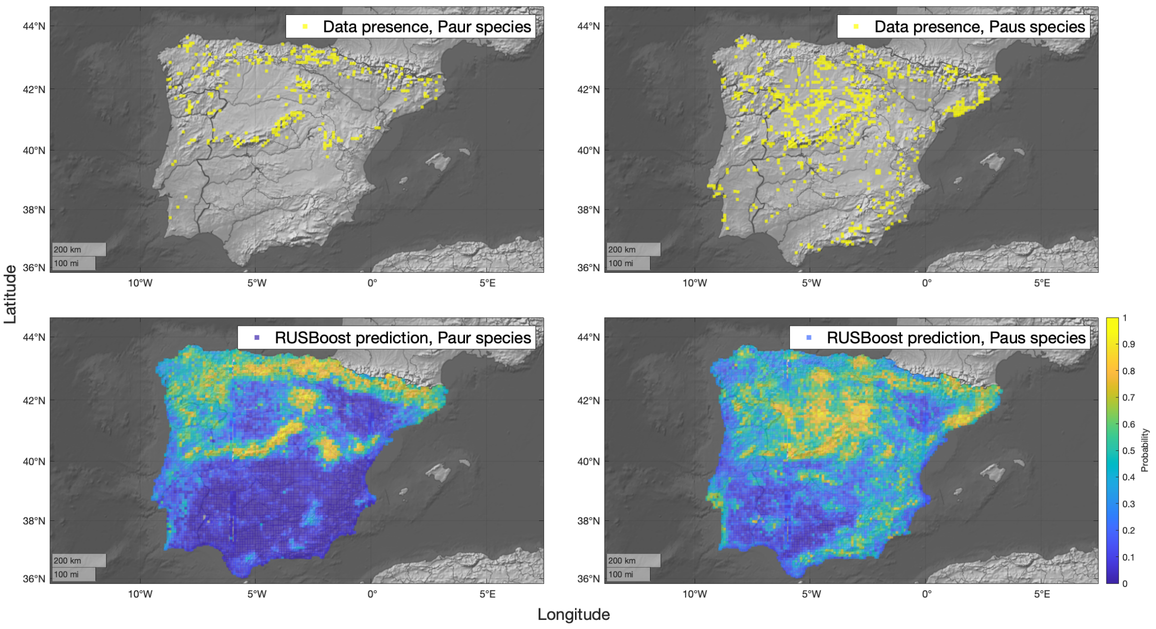
Data presence, Paur species
Data presence, Paus species
RUSBoost prediction, Paur species
RUSBoost prediction, Paus species
44°N
42°N
40°N
38°N
36°N
10°W
5°W
0°
5°E
200 km
100 mi
Latitude
Longitude
Probability
0
0.1
0.2
0.3
0.4
0.5
0.6
0.7
0.8
0.9
1

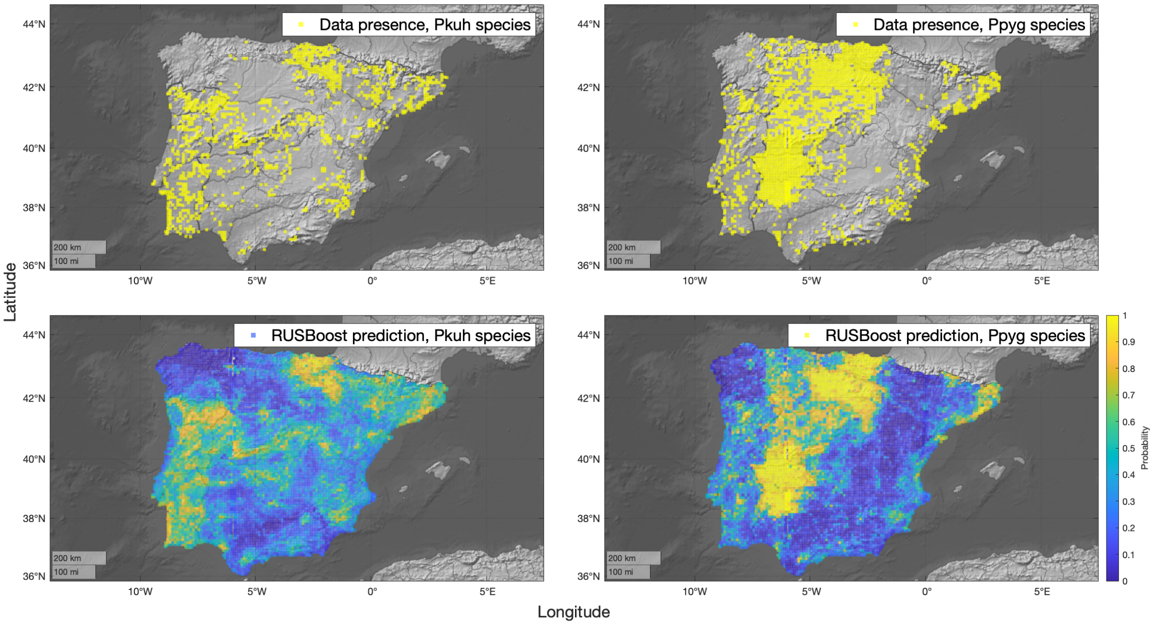
Data presence, Pkuh species
Data presence, Ppyg species
RUSBoost prediction, Pkuh species
RUSBoost prediction, Ppyg species
Latitude
Longitude
Probability
44°N
42°N
40°N
38°N
36°N
10°W
5°W
0°
5°E
200 km
100 mi

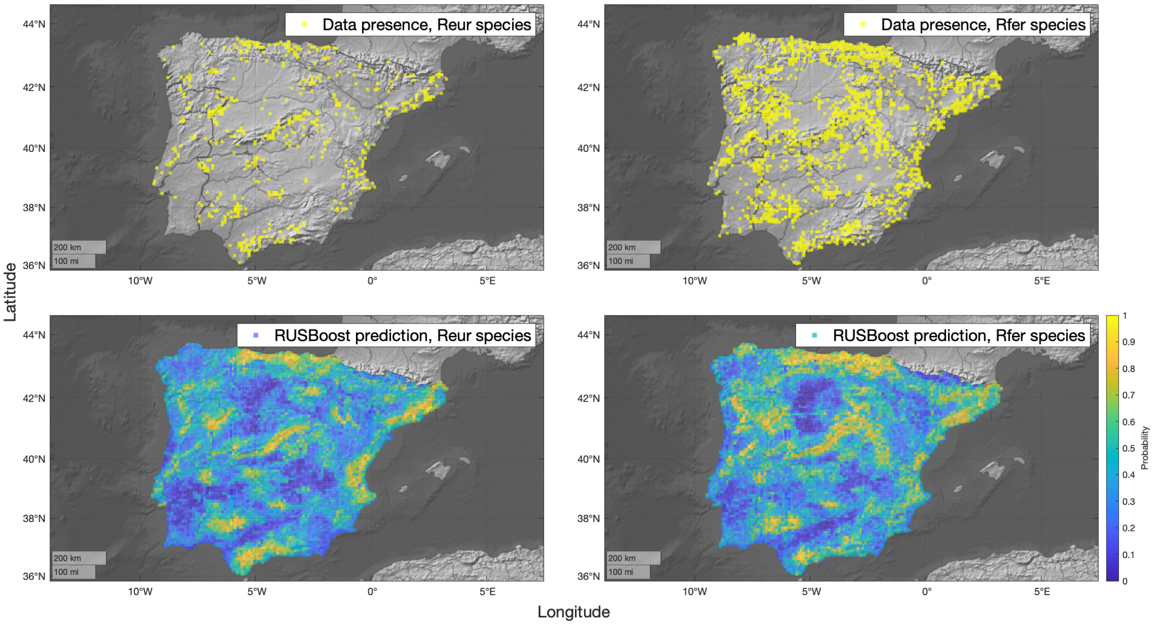
Data presence, Reur species
Data presence, Rfer species
RUSBoost prediction, Reur species
RUSBoost prediction, Rfer species
Latitude
Longitude
Probability
44°N
42°N
40°N
38°N
36°N
10°W
5°W
0°
5°E
200 km
100 mi

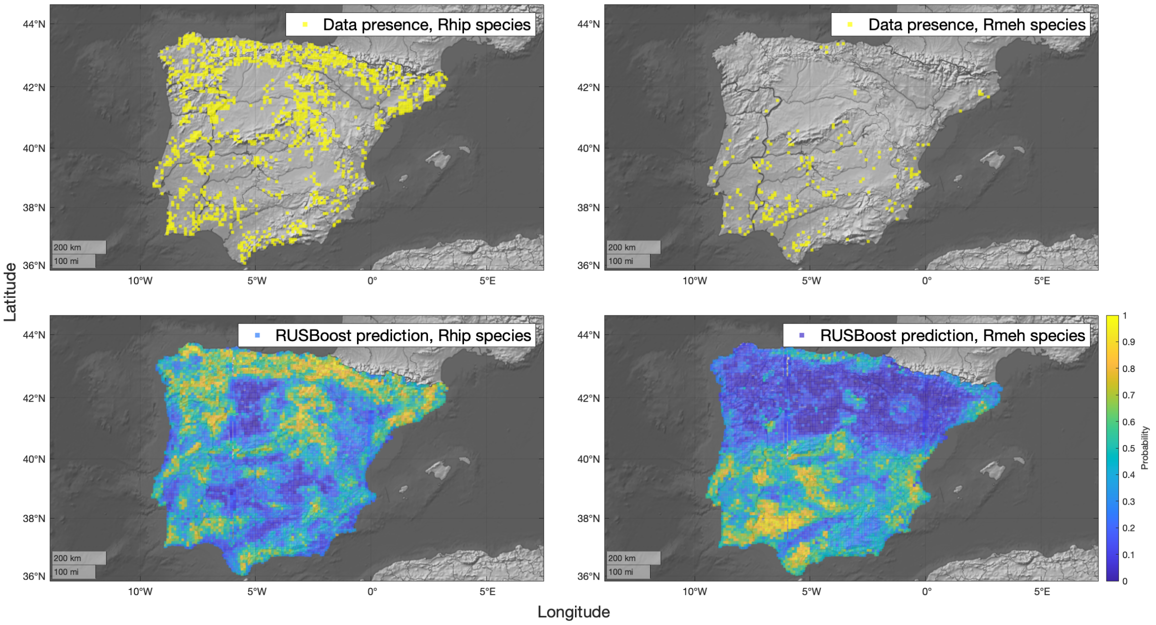
Data presence, Rhip species
Data presence, Rmeh species
RUSBoost prediction, Rhip species
RUSBoost prediction, Rmeh species
44°N
42°N
40°N
38°N
36°N
10°W
5°W
0°
5°E
200 km
100 mi
Latitude
Longitude
Probability
0
0.1
0.2
0.3
0.4
0.5
0.6
0.7
0.8
0.9
1

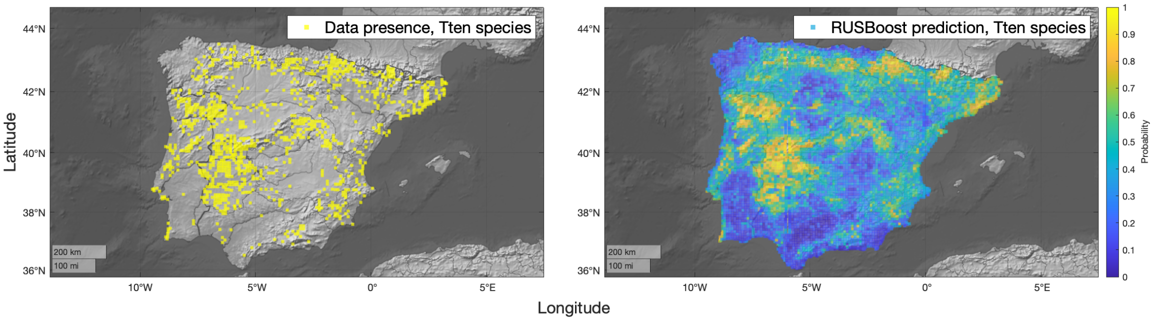
Data presence, Tten species
RUSBoost prediction, Tten species
44°N
42°N
40°N
38°N
36°N
10°W
5°W
0°
5°E
200 km
100 mi
Latitude
Longitude
Probability
0
0.1
0.2
0.3
0.4
0.5
0.6
0.7
0.8
0.9
1
